# FluxMD: High-Throughput Protein Stress Barcoding for Dynamic Structural Analysis

**DOI:** 10.1101/2025.06.17.659767

**Authors:** Jonathan Myunghyun Jeong

## Abstract

Proteins manifest biological functions through precisely orchestrated mechanical responses, yet classical molecular dynamics (MD) faces a profound conundrum: atomistic accuracy is intrinsically antithetical to proteomic scalability. Here we introduce FluxMD, a GPU-accelerated computational framework designed explicitly to surmount this dilemma by translating ligand-protein interactions into unique mechanical fingerprints. FluxMD systematically probes protein surfaces using brief, helically guided Brownian trajectories around rigid receptors, integrating residue-level non-covalent perturbations into a reproducible *stress barcode*. Benchmarking against glutathione peroxidase 4 (GPX4), an essential ferroptosis regulator, exemplifies FluxMD’s predictive sensitivity: the minimalist electrophile TMT10 elicits sharply defined mechanical flux patterns localized at critical nucleophilic residues, whereas the structurally bulky ML162 disperses mechanical signals diffusely. FluxMD enables rapid and mechanically nuanced structural analysis, and is anticipated to adeptly triage biologically salient candidates from vast AlphaFold- and diffusion-generated protein repertoires.

## Introduction

AlphaFold (AF) and diffusion models have *democratized* structural biology, predicting over 200 million protein structures, effectively saturating the UniProt proteome[1, 2]. This structural abundance flips traditional pipelines, placing structure before function, leaving vast archives of *in silico* folds whose biophysical integrity and ligandability remain speculative. Experimental validation, however, cannot scale comparably, highlighting a critical demand for **high-throughput, dynamics-aware filters** to sift biologically relevant structures from merely plausible ones.

Classical molecular dynamics (MD), the *gold standard* for mechanistic insights[3, 4], is prohibitively slow at proteomic scales. All-atom simulations with explicit solvent scale quadratically, consuming weeks of GPU time to access microsecond-level dynamics for even moderate-size systems[5]. Analysis compounds this difficulty; microsecond simulations generate terabytes of data, whose mechanistic interpretation demands laborious manual curation[6, 7]. Enhanced sampling techniques (e.g., metadynamics, accelerated MD) partially mitigate sampling limitations but at exponentially higher computational and expertise cost[8–11]. Thus, MD is poorly suited to rapidly prioritize among millions of predicted protein structures.

Even methods explicitly measuring mechanical stresses face intrinsic challenges. Virial (Cauchy) stress calculations critically depend on assigning residue volumes, an inherently ill-posed problem[12, 13]. Van der Waals volumes underestimate interfacial stress substantially, whereas Voronoi-based methods introduce discontinuities, inflating stress estimates dramatically upon minute structural perturbations[13]. Consequently, identical molecular force fields produce highly inconsistent stresses, severely hindering cross-protein comparisons and obscuring genuine mechanical phenomena, such as allostery[12].

Furthermore, AF’s adoption exposes a deeper issue—the *static-structure paradox*. High-confidence AF predictions (high pLDDT scores) reflect coordinate precision, not dynamical stability[14, 15]. Thus, many structurally accurate models conceal latent dynamics, cryptic cavities, or allosteric instabilities invisible to static inspection[16– 18]. Conventional docking compounds this issue by assuming rigid scaffolds, systematically missing induced-fit or allosteric phenomena[19, 20]. Structural biology thus stands at a paradoxical juncture—rich in static structures yet impoverished in scalable dynamics-informed methodologies[21, 22].

Here, we introduce **FluxMD**, a GPU-accelerated computational platform for high-throughput mechanical screening. FluxMD treats ligand-protein interactions as systematic mechanical probes, orchestrating thousands of brief, helical Brownian trajectories around quasi-rigid receptors (Fig.1). Non-covalent force vectors from these trajectories are distilled into intuitive per-residue *flux metrics*:

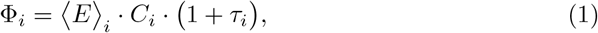

where ⟨*E*⟩*_i_* is mean interaction energy (enthalpy), *C_i_* quantifies directional coherence (directional entropy), and *τ_i_* accounts for temporal variance (magnitudinal entropy). This yields a iteratively reproducible *stress barcode*, clearly delineating mechanically stabilizing from destabilizing residues. FluxMD rapidly screens vast protein libraries, delivering robust mechanical signals far faster than conventional MD.

**Fig. 1.**
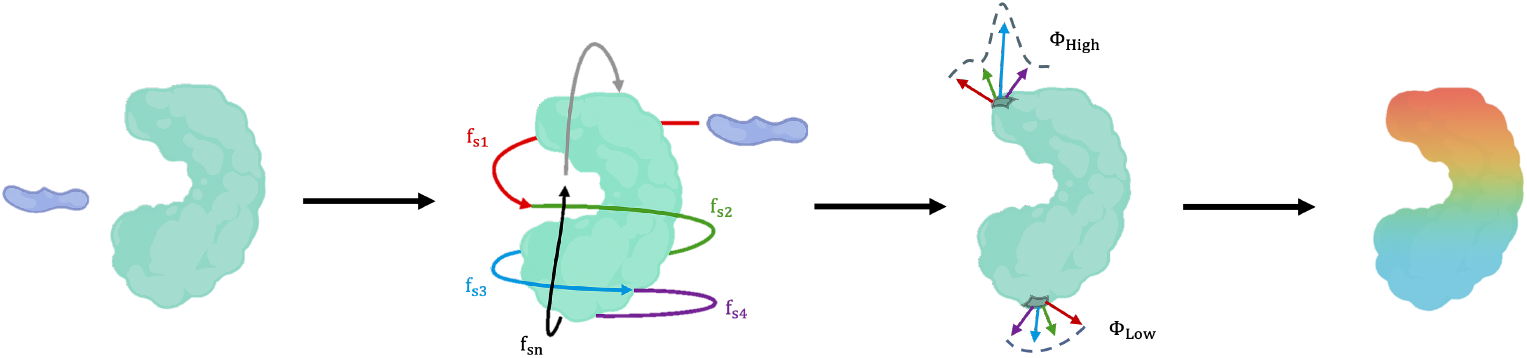
Overview of FluxMD workflow. Ligands sample the receptor surface in controlled helical trajectories (left), generating directional force vectors (center). Aggregation of these vectors produces a spatially resolved stress map, stressing residues subjected to high perturbation (right).

## Results

### Biochemical context

Ferroptosis is an iron-dependent oxidative cell death characterized by lipid peroxide accumulation, implicated in cancer and neurodegenerative diseases [23]. Its primary regulator, glutathione peroxidase 4 (GPX4), detoxifies lipid peroxides, thereby inhibiting ferroptosis. Stockwell *et al.* previously identified GPX4’s active-site selenocysteine (Sec46/U46) and the surface cysteine (Cys66) as nucleophilic residues crucial for covalent inhibition [24]. To assess whether FluxMD’s mechanical analysis independently recapitulates this nucleophilicity profile, we studied GPX4 wild-type (WT) and variants (U46C, U46C/C66S), explicitly focusing on GPX4’s catalytic triad (U46, Q81, and W136; Fig.2b)[25].

**Fig. 2.**
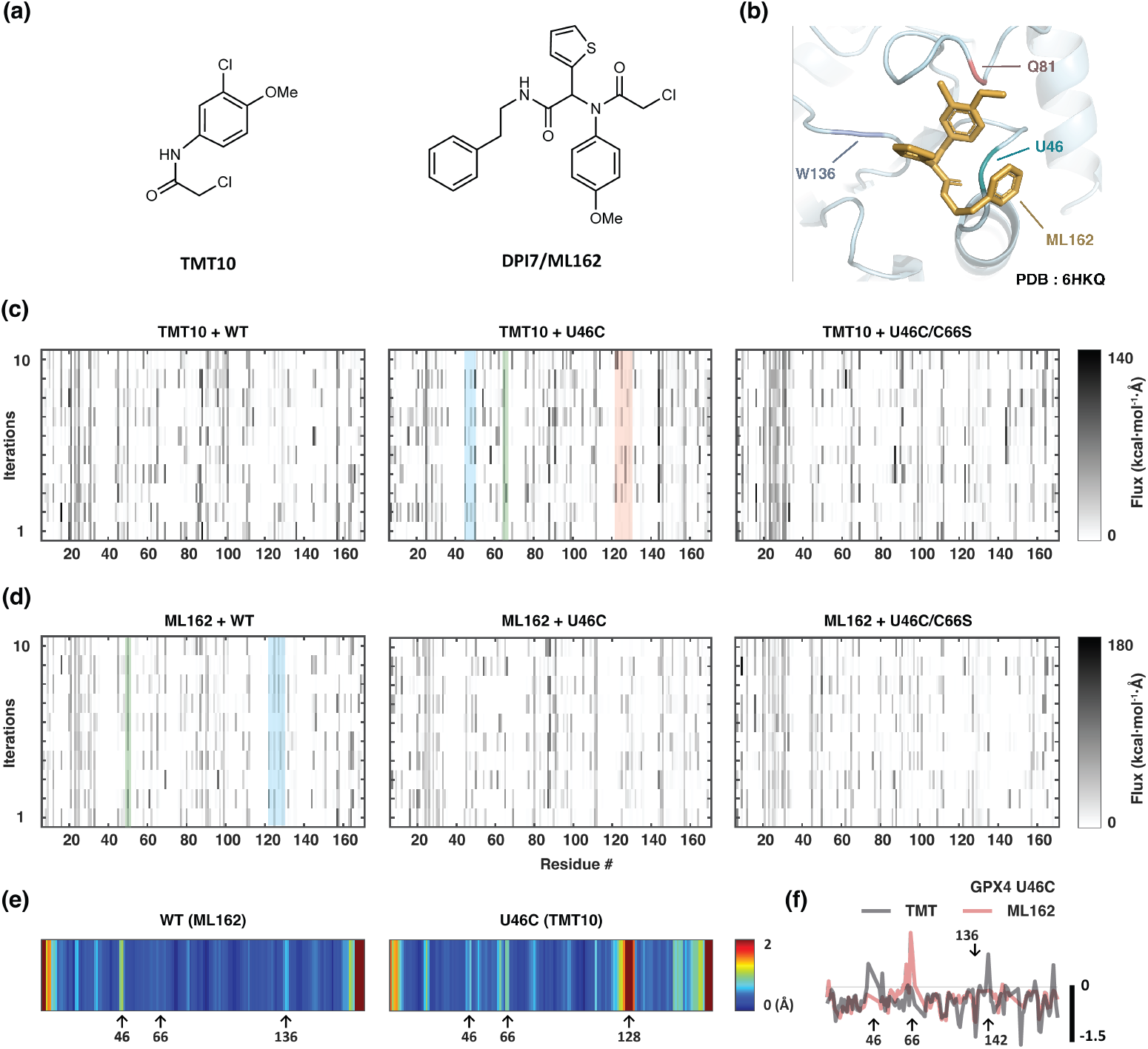
Stress barcode analysis distinguishes ligand-specific mechanical perturbations in GPX4. **(a)** Planar Chemical structures of TMT10 and ML162. **(b)** ML162 binding pose at GPX4 Sec46 (PDB: 6HKQ; WT with surface C66S mutation), highlighting the catalytic triad residues (U46, Q81, W136). **(c, d)** Stress barcode reproducibility across bootstrap replicates (n = 10) for TMT10 (**c**) and ML162 (**d**) binding to GPX4 WT and variants (U46C, U46C/C66S). Inverse grayscale encodes signed flux (kcal·mol^−1^·Å), differentiating stabilizing from destabilizing mechanical interactions. **(e)** RMSD heatmaps illustrate residue-specific structural perturbations: ML162 with WT (left, 6HN3 vs. 6HKQ), and TMT10 with U46C mutant (right, 7L8K vs. 7U4K). **(f)** Gaussian-smoothed flux profiles (*σ* = 2) comparing mechanical resolution at critical residues between TMT10 (gray) and ML162 (red) in GPX4 U46C.

### Barcode sharpness tracks ligand bulk

TMT10 generates sharply defined flux barcodes centered clearly at residue 46 in wild-type GPX4, alongside a thinner, weaker barcode at residue 66 (Fig. 2c). Upon introducing the U46C mutation, flux intensity at residue 46 significantly diminishes, whereas signals at residue 66 remain stable. For ML162, wild-type GPX4 exhibits broad flux density around residue 46 with weaker signals at residue 66 (Fig. 2d). The U46C mutation similarly decreases flux at residue 46 without substantially altering residue 66, although flux intensity generally remains diffuse, reflecting ML162’s larger molecular architecture. The double mutant (U46C/C66S) barcodes largely merge features from both WT and single mutant signatures, reinforcing residue 66’s role as a stabilizing mechanical “stress sink.”

### Orthogonal observables converge

Residue-level RMSD analysis further supports ligand-specific perturbations identified by FluxMD barcodes (Fig. 2e). For WT GPX4 bound to ML162, residues U46 and, less prominently, C66 exhibit moderate structural deviations, consistent with its diffuse barcode signature. U46C bound to TMT10 presents a distinct increase in RMSD primarily at residue C66 and an apparent large deviation at I128. Notably, the RMSD peak at I128 arises is expected due to structural constraints from using PDB:7U4K (U46C-R152H) for comparison, an unavoidable artifact caused by limited ligand-protein complex data. Nevertheless, the RMSD increase at residue 66 reliably supports FluxMD’s barcode results. Additionally, smoothed flux profiles (Fig. 2f) confirm residue-specific mechanical perturbations, notably a clear divergence between the minimalist electrophile TMT10 and the structurally bulkier ML162. Thus, independent structural measures reinforce FluxMD’s capacity to mechanically resolve nucleophilic residues consistent with biochemical nucleophilicity mapping by Stockwell *et al.*

## Discussion

FluxMD transforms biomolecular dynamics into tractable mechanical fingerprints. Unlike existing methods such as Perturbation Response Scanning (PRS)[26], which applies internal forces to map allostery, or FTMap[27], which statically scans protein surfaces, FluxMD captures mechanical responses externally through ligand trajectories, avoiding normal-mode or explicit solvent calculations. Traditional MD’s explicit solvent and microsecond demands become unnecessary as FluxMD distills high-dimensional dynamics into concise residue-level flux signatures.

FluxMD relies on three innovations: first, a flux metric Φ*_i_* = ⟨*E*⟩*_i_ × C_i_ ×* (1 + *τ_i_*) unifying energetic and kinetic signals into one quantitative measure; second, a volume-independent approach resolving longstanding problems with virial stress analyses; third, GPU-native operations enabling unprecedented proteomic throughput. This approach achieves efficient allosteric mapping comparable to PRS but via fundamentally different external probing.

The GPX4 case illustrates FluxMD’s diagnostic precision, linking ligand bulk directly to mechanical resolution. The minimalist TMT10 electrophile generated sharply defined flux peaks at nucleophilic hotspots, clearly differentiating sites of covalent modification. In contrast, bulkier ML162 induced diffuse flux patterns, dispersing mechanical signals due to steric hindrance. Hence, compact ligand structures maximize FluxMD’s mechanical clarity.

FluxMD’s simplifications—implicit solvation, rigid receptors, nanosecond sampling—are deliberate, prioritizing practical throughput over atomic-resolution complexity. This design makes FluxMD uniquely suited to screening vast AlphaFold-derived libraries, quickly identifying mechanically stable candidates worthy of experimental effort.

The reproducibility of FluxMD’s stress barcode converts subjective structural evaluation into systematic, quantitative fingerprinting, establishing it as a quality-control method within protein prediction pipelines. However, absolute flux intensities and exact hotspot localization might vary across different force fields, necessitating future systematic benchmarking. Current limitations, including uniform temporal weighting (*τ*) and omission of explicit solvent effects, might misrepresent certain dynamic phenomena. Incorporating explicit solvent dynamics and frequency-weighted *τ_i_* terms calibrated experimentally will enhance FluxMD’s physiological accuracy.

Interpretation of the GPX4 double mutant (U46C/C66S) remains cautious. The benchmarked study[24] employed MALDI-MS under denaturing conditions, leaving unclear whether the mutants’ lack of covalent modification arises from genuine structural immunity or compromised folding. FluxMD’s mechanical profiling could potentially distinguish these scenarios by identifying either stable, non-reactive mutants or structurally compromised variants through distinct mechanical signatures, awaiting experimental validation.

In conclusion, FluxMD provides an efficient, scalable framework for dynamically validating protein structures, rapidly linking mechanical insights to biological function. By balancing simplicity and rigor, it democratizes mechanical profiling, accelerating the translation of computational predictions into actionable biochemical insights.

## Methods

FluxMD converts ligand binding into an external mechanical assay, running via either CPU (standard) or GPU (zero-copy UMA) execution paths. Both paths follow the same logic: (i) trajectory generation, (ii) force evaluation via Rosetta REF15, and (iii) statistical reduction into residue-level flux vectors. Complete source code and installation instructions are available at the public GitHub repository (https://github.com/myunghyunj/FluxMD-simulation).

### Protein and ligand preparation

#### Wild-type GPX4 preparation

Wild-type GPX4 (PDB: 6HN3) includes selenocysteine (Sec46), unsupported by FluxMD’s REF15 implementation. Sec46 was thus computationally replaced by cysteine (U46→C46) using Biopython 1.85, preserving backbone coordinates exactly (RMSD = 0.00 Å).

#### Variant GPX4 preparation

The GPX4-U46C variant was obtained directly from PDB:7L8K [28]. GPX4-U46C/C66S was modeled using AlphaFold3 (seed: 239214584, pTM=0.91).

#### Ligand (SMILES → PDB) conversion

Ligands used:

- TMT10: `C1(=CC(=CC=C1OC)N(C(CCl)=O)[H])Cl`
- ML162: `[s]1c(ccc1)C(N(c3cc(c(cc3)OC)Cl)C(=O)CCl)C(=O)NCCc2ccccc2`

Conversion from SMILES to PDB was done primarily using the NCI CACTUS webserver. Due to CACTUS occasionally omitting aromatic ring connectivity, FluxMD automatically reconstructs missing `CONECT`records. If CACTUS fails, OpenBabel 3.1.1 is used. Conversion details are documented on GitHub under “`SMILES`→`PDB Converter`“.

### Force-field Implementation and Energy Model

FluxMD employs an internal implementation of the Rosetta REF15 energy function, evaluating forces using standard terms (fa atr, fa rep, fa sol, fa elec, hbond). The model uses a sigmoidal distance-dependent dielectric that transitions from =10 at close range to =80 at larger distances, providing a physically realistic representation of electrostatic screening. Atom typing follows Rosetta’s 167-type convention; missing types default to carbon with warnings emitted. Protonation states are dynamically assigned via the Henderson–Hasselbalch equation at a physiological condiction (pH and temperature).

### Trajectory engine

#### Adaptive helical orbits

Ligands explore protein surfaces via overdamped Langevin dynamics with a timestep of 40 fs. This interval deliberately exceeds conventional MD steps (1–2 fs) by ignoring intramolecular vibrations above 25 THz (e.g., bond stretches). Given our rigid-body protein assumption—where highly stressed residues indicate potential sites of conformational change—this choice rigorously satisfies the Shannon–Nyquist criterion for biologically relevant intermolecular motions (0–10 THz). Principal Component Analysis (PCA) determines adaptive trajectory shapes: spherical cocoon trajectories for globular proteins and cylindrical (helical) trajectories for elongated molecules. Each translational step includes rotations through 36 evenly spaced Euler angles, generating ∼ 10^5^ discrete ligand poses per run, optimizing both computational efficiency and critical interaction resolution.

#### Collision handling and corrections

When heavy-atom pairs violate van der Waals radii (0.8 × (*r*_vdw,1_ + *r*_vdw,2_)), the step is rejected. Corrective maneuvers include radial retreat, incremental trajectory adjustments, or larger jumps to escape sterically hindered regions.

#### Multi-layer cocoon schedule

Trajectories are generated in concentric layers at progressively decreasing mean distances (default: 20–50 Å) from the protein surface. Ligand–protein distances fluctuate via momentum, Brownian jitter, and periodic close approaches (minimum 1.5–2.0 Å) to robustly sample short-range interactions.

#### Rigid receptor rationale

Protein backbone and sidechains remain fixed, isolating rapid surface perturbations from slower conformational drifts. To characterize internal strain (stress), internal protein energetics are precomputed once using REF15, enhancing computational efficiency and enabling full GPU optimization without loss of structural context.

### Flux metric

The residue-level mechanical flux (Φ*_i_*) is defined as:

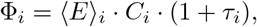

where ⟨*E*⟩*_i_* (kcal·mol^−1^) is the residue’s mean signed interaction energy, *C_i_* ∈ [0, 1] quantifies directional coherence defined as

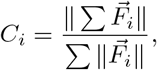

and *τ_i_* is the temporal coefficient of variation

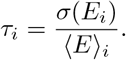

Negative flux values indicate net attractive interactions (stabilizing), whereas positive values reflect net repulsion (destabilizing).

### Statistical validation

Statistical significance of residue-level flux (Φ) is evaluated by two complementary methods, depending on computational workflow.

#### Bootstrap percentile test (CPU workflow)

The standard CPU pipeline employs non-parametric bootstrap resampling (10^3^ iterations). Flux values are resampled with replacement to empirically estimate two-tailed *p*-values, calculated as twice the fraction of bootstrap means more extreme than the observed mean flux.

#### Student’s t-test (UMA/GPU workflow)

The UMA pipeline and recovery scripts employ a parametric one-sample Student’s *t*-test (SciPy ttest 1samp), assessing the null hypothesis (Φ = 0):

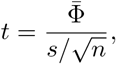

where 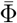 denotes mean flux, *s* standard deviation, and *n* iteration count. Resulting two-tailed *p*-values quantify deviation significance from zero.

Each FluxMD iteration (or run) comprises ten independent cocoon trajectories (*N* = 10). Residues exhibiting flux magnitudes within the top decile and significance at *p <* 0.05 are classified as stress hotspots. However, for visual clarity in Fig. 2, these hotspots are not individually marked, as the analysis revealed them to be highly concentrated on either N- and C-termini of the respective protein’s residue that finely collapses to the RMSD difference distribution aspects, thus regarded as artifacts.

### Data visualisation and recovery

Barcodes and flux profiles are rendered with Matlab (Mathworks®, MA, USA) and elaborated in Adobe Illustrator (Adobe Inc., CA, USA). All trajectory parameters, energy caps and reproducibility scripts are available in the open-source FluxMD repository.

## Acknowledgements

The author acknowledges that computational analyses were conducted using personally available computing resources, due to the absence of dedicated external funding or institutional computational infrastructure support.

FluxMD was developed independently, building upon legacy work established during my previous term project (Autumn 2023) on protein folding analysis (https://github.com/jaehee831/protein-folding-analysis-bioinformatics). I thank my teammates from the 2023 project for discussion feedback that significantly enriched the theoretical foundation of this work.

## Author Contributions

JMJ solely conceptualized, coded, and wrote this paper.

## Data Availability

FluxMD v1.3.0 is accessible at https://github.com/myunghyunj/FluxMD-simulation/ under the MIT license. Relevant raw data presented in this paper is also available in the repository.

## Supplementary Information

Supplementary data and additional analyses supporting this manuscript are available upon request. Raw data are all available in GitHub.

